# Critical thermal maxima and oxygen uptake in *Elysia viridis* (Montagu, 1804), a sea slug capable of photosynthesis

**DOI:** 10.1101/2023.06.19.545621

**Authors:** Elise M. J. Laetz, Can Kahyaoglu, Natascha M. Borgstein, Michiel Merkx, Sancia E. T. van der Meij, Wilco C. E. P. Verberk

## Abstract

Photosynthetic animals produce oxygen internally, providing an ideal lens for studying how oxygen dynamics influence thermal sensitivity. The sea slug, *Elysia viridis*, can retain functional chloroplasts from its food alga *Bryopsis plumosa* for months, but retention is limited when fed *Chaetomorpha* sp., limiting potential oxygenic benefits. We fed slugs each alga and exposed them to 17°C (their current yearly maximum temperature) and 22°C (the increase predicted for 2100), to examine plasticity in thermal tolerance and changes in oxygen uptake when fed and starving. We also examined slugs under increased illumination to examine a potential tradeoff between increased oxygen production, and a faster rate of chloroplast degradation. Following exposure to these conditions, we performed ramping trials, subjecting them to acute thermal stress to determine their thermal tolerance. We also measured oxygen uptake before and after ramping. We observed increases in thermal tolerance for specimens exposed to 22°C, indicating they acclimated to temperatures higher than they naturally experience. Fed slugs exhibited higher rates of oxygen consumption before exposure to acute thermal stress, and suppressed their oxygen uptake more after it, than starved slugs. Under higher light, slugs exhibited improved thermal tolerance, possibly because increased oxygen production alleviated host oxygen limitation. Accordingly, this advantage disappeared later in the starvation period when photosynthesis ceased due to chloroplast digestion. In conclusion, *E. viridis* can suppress metabolism to cope with heat waves, however, starvation influences a slug’s thermal tolerance and oxygen uptake, so continuous access to algal food for chloroplast retention is critical when facing thermal stress.

**Summary Statement:** Oxygen has been implicated in determining an ectotherm’s thermal sensitivity. Examining photosynthetic (and therefore oxygen-producing) sea slugs under various conditions helps elucidate how oxygen and other factors impact thermal tolerance.

## Introduction

Temperature and oxygen availability are perhaps the most critical abiotic factors for multicellular life and both have pervasive effects across all levels of biological organization. Oxygen availability is limited in aquatic habitats because oxygen diffuses ∼300,000 times slower in water than in air (Dejours, 1981). Temperature modulates the availability of oxygen by changing the solubility and diffusivity of oxygen in water and it also changes the thickness of boundary layers via the effects on the viscosity of water (Graham, 1990; Verberk *et al*., 2011; Verberk and Atkinson, 2013). Rising global temperatures due to climate change can deoxygenate aquatic habitats because oxygen solubility in water decreases with elevated temperatures, while metabolic rates increase, which in turn increases the demand for oxygen to maintain aerobic respiration. Given these links between oxygen dynamics and temperature, hyperthermal- and hypoxic stress can amplify one another (Deutsch *et al*., 2015; Verberk, Durance, *et al*., 2016).

Exposure and sensitivity to heat vary across aquatic and terrestrial realms (Pinsky *et al*., 2019), and ectotherms inhabiting the intertidal and uppermost subtidal zones are particularly exposed to heat stress (Pörtner, 2001; Cereja, 2020). The maximum temperatures present in intertidal habitats are strongly associated with the upper thermal limits observed in their ectotherm inhabitants (Stillman and Somero, 2000; Stillman, 2003), although these limits and the effects of thermal stress vary by taxa and latitude (Vinagre *et al*., 2019). In other words, many ectotherms already experience conditions that approach the tolerance limits that define their thermal windows (Vinagre *et al*., 2019; Cereja, 2020).

Critical thermal maxima are often investigated by monitoring various biological changes (such as running performance or metabolic rate/oxygen uptake) in individuals exposed to increasing temperatures (Clusella-Trullas *et al*., 2011; Vasseur *et al*., 2014). An animal’s upper thermal limit is usually defined as the critical thermal maximum at which it loses voluntary muscle control and is no longer able to try to escape from conditions that will result in its death (Cowles and Bogert, 1944) or it enters a heat-induced coma (Lutterschmidt and Hutchison, 1997; Armstrong *et al*., 2019). Although critical thermal maxima are often portrayed as being fixed properties of a species, they can be modulated via acclimation and heat hardening. In the case of thermal acclimation, exposure to mild increases in temperature result in physiological changes that allow an individual to better tolerate hyperthermal stress (e.g. Gunderson and Stillman, 2015). In the case of heat hardening, a short exposure to intense heat triggers a heat-shock response, which leads to a reduction of energy costs associated with protein synthesis and ion pumping (Feder and Hofmann, 1999). These energetic reductions can further improve survival of individuals subjected to subsequent acute heat stress (Verberk and Calosi, 2012), but can have long-term fitness costs (Krebs and Loeschcke, 1995).

Since climate change is implicated in both rising average temperatures and the increase in extreme weather events such as heat waves, the effects of both chronic warming and acute heat-shocks are relevant when trying to predict whether or not species can cope with climate change. Although the energetic consequences of exposure to temperatures outside an ectotherm’s thermal window are increasingly understood, the causes underpinning thermal limits in ectotherms remain unclear. Thermal effects on both oxygen availability and demand have been used to argue that insufficient oxygen may be a key factor in defining an ectotherm’s thermal window (Hochachka and Somero, 2002; Pörtner and Farrell, 2008; Verberk, Overgaard, *et al*., 2016; Hoefnagel and Verberk, 2017). However, the generality and testability of this oxygen limitation hypothesis are debated (summarized in Verberk, Overgaard, *et al*., 2016; Jutfelt *et al*., 2018; Pörtner *et al*., 2018). Surveying a wide range of species from different clades and studying model organisms with unique physiological attributes can help to disentangle specific aspects of the relationships between temperature, oxygen demand and availability.

Animals that share a symbiotic relationship with photosynthetic organisms could provide a lens with which to study the interactions between temperature, oxygen demand and oxygen availability because the photosynthesis occurring in the symbionts produces oxygen that can be used to sustain the animal’s aerobic respiration and even oxygenate the environment. Some species of sea slugs (Gastropoda: Sacoglossa) stand out as ideal study subjects for such experiments because they steal and retain functional chloroplasts from their algal food sources in a process called functional kleptoplasty (Rumpho *et al*., 2010). Once incorporated in the slug, these kleptoplasts can continue to function, sometimes for months, producing both oxygen and photosynthates such as carbohydrates (Trench *et al*., 1973; Hinde and Smith, 1975; Laetz *et al*., 2016; Laetz and Wägele, 2018) or lipids (Pelletreau *et al*., 2014). These photosynthates can provide nutrition to starving slugs during periods of food unavailability (due to the pelagic stages of some algal life cycles) or food inaccessibility (due to calcification of the algal thallus that prevents slugs from feeding) (Marin and Ros, 1992). How much nutrition is provided by photosynthesis is debated (Cartaxana *et al*., 2017; Rauch *et al*., 2017; Laetz and Wägele, 2018) and highly dependent on the algal and slug species/populations involved (Wägele and Martin, 2014). However, kleptoplasty does not produce enough photosynthates to fully support a starving slug, i.e. the slug is not photoautotrophic (Hinde and Smith, 1975; Cartaxana *et al*., 2017; Rauch *et al*., 2018).

The emerald sea slug, *Elysia viridis* (Montagu, 1804) has been a model species for understanding functional kleptoplasty for over 50 years (Taylor, 1968). It inhabits the uppermost subtidal zone from the Mediterranean Sea to the British Isles and the northeastern Atlantic coast of Scandinavia (Baumgartner *et al*., 2015; Laetz *et al*., 2016). *Elysia viridis* is known to feed on a multiple, chlorophyte genera including, *Codium* Stackhouse, 1797 *, Bryopsis* Lamouroux, 1809 *, Chaetomorpha* Kützing, 1845 *, Cladophora* Kützing, 1843, and the rhodophyte genus, *Griffithsia* Agardh, 1817, (Händeler and Wägele, 2007; Baumgartner *et al*., 2015; Rauch *et al*., 2018). The amount of time it can retain functional chloroplasts ranges from 1-2 months when fed *Codium spp.* and *Bryopsis spp.*, down to a few weeks when fed *Chaetomorpha spp.* (Trowbridge *et al*., 2008; Rauch *et al*., 2018). Chloroplast retention has not been observed in *E. viridis* feeding on *Cladophora spp.* or *Griffithsia spp*. Since oxygen is produced as a byproduct of photosynthesis within the slug’s body, it is available for aerobic respiration. Internal oxygen production could be especially important when demand is high, such as in warm waters, but this remains uninvestigated.

In this study, we investigated the critical thermal maxima for *E. viridis* specimens under varying biotic and abiotic conditions to examine how environmental stress affects solar-powered slug heat tolerance. We also measured their oxygen uptake rates before and after assessing their critical thermal maxima to determine how their energy metabolism changed after exposure to acute heat stress. Since photosynthetic activity decreases during starvation due to chloroplast digestion (Laetz *et al*., 2016; Frankenbach *et al*., 2021), and periods of food unavailability have been observed for this species (Baumgartner, 2014), we measured heat tolerance and metabolism at different points during a starvation period.

In order to make inferences about how these slugs will tolerate prolonged temperatures above those in which they naturally occur, we compared slugs exposed to 17°C (daily mean maximum summer temperature in 2020 below the thermocline), to slugs we exposed to 22°C, reflecting the 5°C increase in North Sea surface temperatures predicted by a climate model for the year 2100 (Klein Tank and Lenderink, 2009) (Experiment 1). In this experiment, we fed *E. viridis* either *Bryopsis plumosa* (Hudson) C. Agardh, 1823 or *Chaetomorpha* cf. *linum* (O.F. Müller) Kützing, 1845. *Elysia viridis* can retain chloroplasts from other algal species within each of these genera, but these algal species have not been assessed. When fed *Bryopsis hypnoides* J.V. Lamouroux, 1809, stolen chloroplasts can remain functional for up to 21 days (Rauch *et al*., 2018). Slugs have been observed feeding on *Chaetomorpha melagonium* (F. Weber & D.Mohr) Kützing, 1845 (Baumgartner and Toth, 2014) but the duration chloroplasts remain active has not been assessed in this species and some authors doubt whether chloroplasts from *Chaetomorpha* sp. can be retained at all (Clark *et al*., 1990; Trowbridge and Todd, 2001). By examining slugs fed these two algal species, we were able to assess how differences in chloroplast retention time and nutritive quality affected thermal tolerance and oxygen uptake.

We also exposed some slugs that were maintained at 17°C and fed *B. plumosa* to a higher light intensity (Experiment 2). The higher light level is expected to increase photosynthetic capacity and oxygen production, at least on the short-term, but could, in the long-term, also lead to an increased breakdown of incorporated chloroplasts due to eventual photodamage and the inability of a slug to repair its chloroplasts (de Vries *et al*., 2013; Christa *et al*., 2018). We identified and present here, environmental factors that underpin a solar-powered slug’s response to acute and prolonged exposure to hyperthermal stress and our predictions for its ability to handle the warmer, deoxygenated ocean of the future.

## Materials and Methods

### Specimen acquisition

*Elysia viridis* specimens were collected from a shallow reef at Bommenede in Zeeland, the Netherlands (51°43’47.5"N 3°58’35.3"E) at a depth of 20cm – 50cm from July, 2020 to February, 2021 (Fig. 1A). Rocks covered in *B. plumosa* were also collected (Fig. 1B). *Chaetomorpha* cf. *linum* was donated by Frits Kuiper Groningen, a local aquarium store.

**Figure 1.**
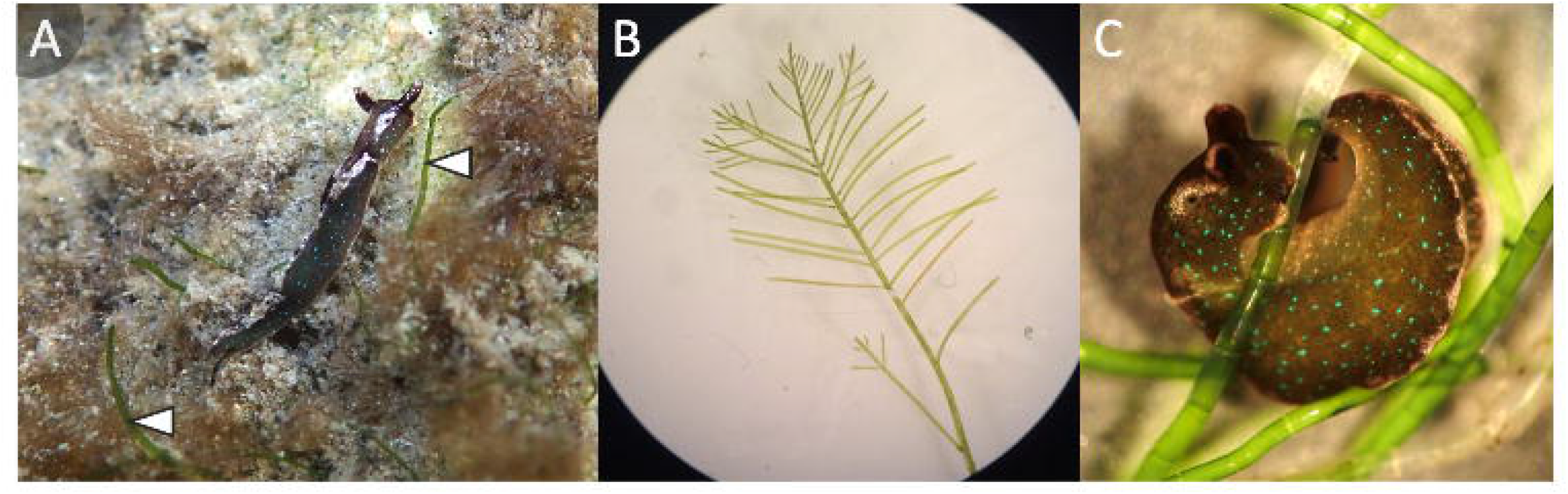
Elysia viridis and its food algae. A) *E. viridis* and small *B. plumosa* thalli (indicated by white arrowheads) under natural conditions at the collection site. *E. viridis* specimen length ∼7mm. B) *Bryopsis plumosa* thallus growth after grazers, primarily *E. viridis*, were removed. Each algal thallus grew to ∼7cm in length and the specimen pictured is ∼5cm in length. C) An *E. viridis* specimen feeding on *C.* cf*. linum.* Specimen length ∼15mm.

Above the thermocline, the temperature reached 23°C in July, 2020, the warmest month at this collection site in 2020 (Rijkswaterstaat, 2020). During July, *E. viridis* migrated below the thermocline though to ∼1.5m depth where they were exposed to a maximum of 17°C, so 17°C was considered the maximum temperature under present conditions (without the +5°C predicted increase due to global warming). All specimens were transported to the laboratory at the University of Groningen for experimentation since cultivation in the laboratory under controlled conditions is not possible for this species.

### Initial acclimation to lab conditions

All slugs were allowed to adjust to lab conditions for a minimum of two weeks before experimentation began. During this time, all slugs were split up into multiple glass tanks with 10 individuals per tank and provided a continuous supply of *B. plumosa,* on which they were observed feeding. Throughout the entire acclimation and experimental periods, partial water changes (∼50%) were performed weekly, providing specimens with fresh, artificial sea water (Instant Ocean - Spectrum Brands, Inc). Each tank was scrubbed every month and given a complete water change to prevent biofilm and/or feces accumulation. Tanks were also outfitted with aerators to ensure the water was mixed and that O_2_ was consistently at full saturation. Each aquarium was provided with 100 µmol m^-2^ s^-1^ full spectrum light (Superfish SLIM LED 45) and the temperature of the climate chamber (Clima Temperatur Systeme T600) where they were housed was maintained at 17°C to match the maximum yearly temperature to which they were exposed in 2020. An overview of the acclimation phase, designs for Experiment 1 and 2 and the treatments used in each experiment can be found in Fig 2.

**Figure 2.**
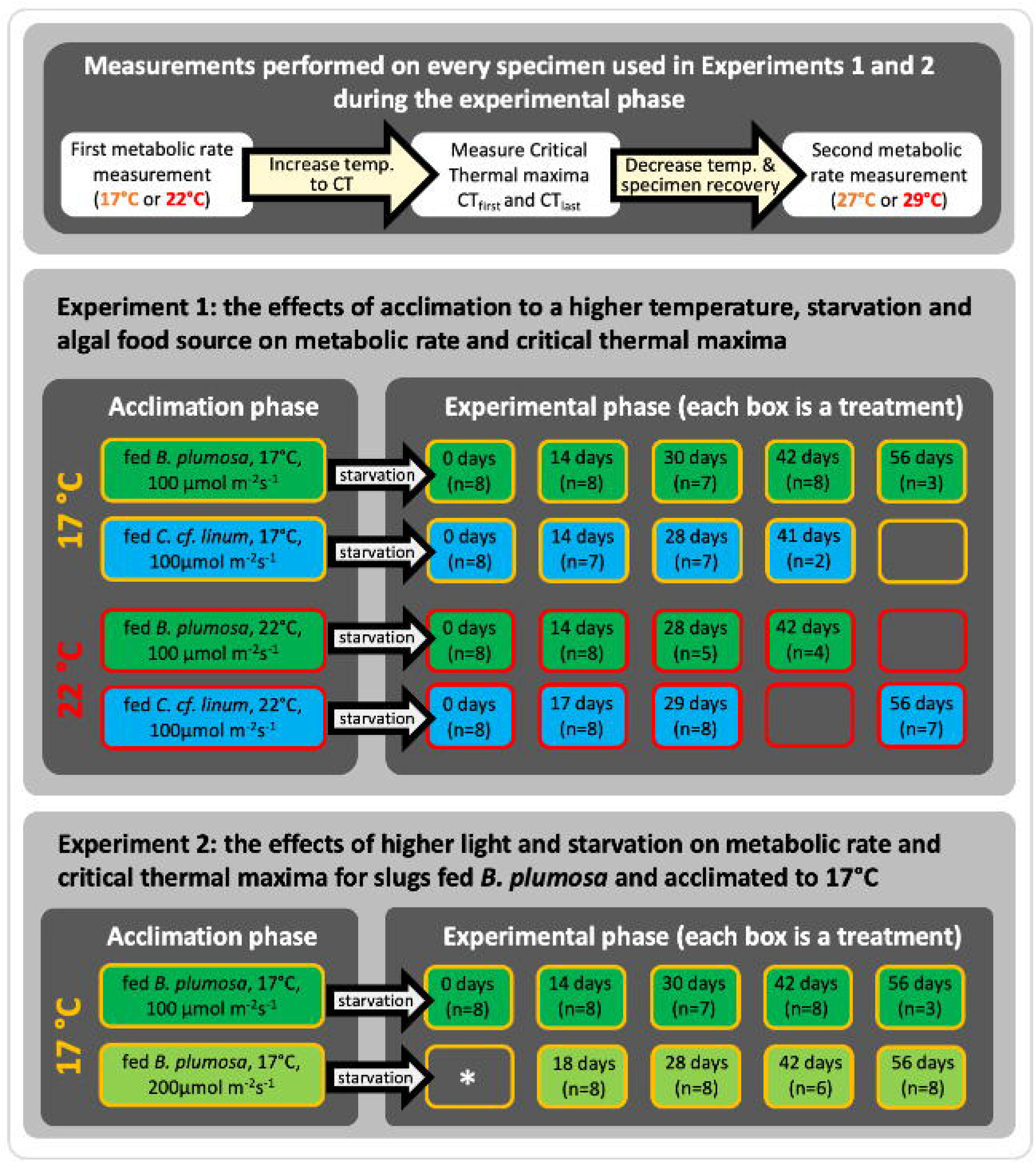
Overview of the design and sampling points for Experiments 1 and 2. Every slug in every treatment was measured according to the same procedure. First, its oxygen uptake (metabolic rate) was measured. Then its critical thermal maxima (CT_first_ and CT_last_) were measured and its oxygen uptake was measured again (top panel). Specimens were fed and exposed to laboratory conditions during the “Acclimation phase”, depicted in the panels on the left. The temperature and light intensity were maintained when specimens were removed from the algae and placed in clean tanks for starvation and experimentation during the “Experimental phase” (right panels). Treatments that are labeled “0 days starved” include specimens that were measured right after the acclimation phase and had therefore fed *ad libitum* until a few hours before experimentation. The number of individuals (n) tested at each sampling point is written below the corresponding sampling point. Empty boxes indicate sampling points we intended to take but weren’t able to take due to specimen mortality or logistical reasons (*).

### Pre-treatments for Experiment 1 - increasing temperatures and a different algal food source

To examine the effects of prolonged exposure to increased temperatures, a change in algal food source, and starvation on oxygen uptake and critical thermal maxima (Experiment 1), approximately 100 slugs were randomly selected and moved to another climate chamber (Clima Temperatur Systeme T600). In this new climate chamber, the temperature was gradually increased to 22°C, at a rate of 1°C per day to ensure that all specimens were slowly exposed to the warmer temperature, which reflects the 5°C temperature increase predicted for the North Sea by 2100 due to global warming (Klein

Tank and Lenderink, 2009). Slugs were then given two weeks to acclimate and feed on *B. plumosa* upon reaching 22°C. After the change in temperature, half of the specimens maintained under each temperature regime (17°C and 22°C) were switched to a diet of *C.* cf*. linum*. The diet switch was done three weeks prior to the start of the experiment to ensure they no longer contained *B. plumosa* chloroplasts (Fig. 1C), as recommended by Frankenbach *et al*. (2021). Following these periods where slugs were exposed to each temperature and diet, each specimen was placed in a clean tank without algae and it entered the experimental phase (described below).

### Pre-treatments for Experiment 2 - increasing the light intensity

In a second experiment, we examined the effect of light intensity on a subset of our specimens (Fig 2). Specimens that were exposed to 17°C and fed *B. plumosa* (as described above) were provided either 100µmol m^-2^ s^-1^ or 200µmol m^-2^ s^-1^ of full spectrum light (Superfish SLIM LED 45) for 12 hours each day. These light intensities were chosen for a few reasons. Natural light intensity varies considerably and is influenced by a multitude of environmental factors (solar exposure, time of year, shade by other objects in the habitat, etc.). Algae can adjust their photoacclimation status to modulate light harvesting under these naturally fluctuating conditions, but this requires nuclear transcription and regulation. *Elysia viridis* lacks algal nuclei, meaning it lacks the ability to photoacclimate and repair/replace many photosystem components when they get damaged by excessive light (Rauch *et al.,* 2015). This means that *E. viridis* hosts kleptoplasts whose photoacclimation status was determined by the light intensities to which the algae they consumed were exposed and themselves photoacclimated (Vieira *et al*., 2009), which occurred during the acclimation phase (i.e. 100 or 200 µmol m^-2^ s^-1^) in our study.

Vieira *et al*. (2009) demonstrated that exposure to 140µmol m^-2^ s^-1^ markedly decreased photosystem II activity in *E. viridis*, shortening the time kleptoplasts remained functional when compared to specimens exposed to 30µmol m^-2^ s^-1^, although photosynthetic efficiency decreased in both light treatments. We chose 100µmol m^-2^ s^-1^ as our “normal” light condition because 30µmol m^-2^ s^-1^ is far lower than these species would regularly experience in the field (which often exceeds 100µmol m^-2^ s^-1^ on a shallow reef).

We chose 200µmol m^-2^ s^-1^ as our high-light treatment because it exceeds the amount of light previously reported to markedly decrease photosystem II activity. Light intensity was measured with an SQ-500 full-spectrum quantum sensor (Apogee Instruments, Inc, USA) using corrections for underwater measurements as instructed by the manufacturer. The quantum sensor was placed underwater in each seawater-filled aquarium and the water bath where each experiment was conducted (described in Experimental phase - calculating the rate of oxygen uptake below) and the light intensity measurements in µmol m-2s-1 were recorded. After this acclimation phase in which the slugs were exposed to either 100µmol m^-2^ s^-1^ or 200µmol m^-2^ s^-1^ for a minimum of two weeks, the slugs were transitioned to the experimental phase (detailed below).

### Experimental phase - measuring the rate of oxygen uptake

The oxygen uptake for each specimen in Experiments 1 and 2 was determined before and after its critical thermal maxima (described in next section) were measured. Oxygen uptake rates were measured via a closed-system respirometry with a Fibox 3 LCD Trace oxygen meter and PSt3 optical sensors that were glued into glass vials (Presens GmBH, Germany). Prior to the measurements, we calibrated the O_2_ sensor spot using a 2- point calibration (100% O_2_ saturated seawater and 0% oxygen: nitrogen gas). The values we report represent net rates of oxygen uptake and are an underestimation of the total oxygen needed to support aerobic respiration. This is because each specimen also produced oxygen via photosynthesis. We did not quantify the relative contribution of oxygen produced via photosynthesis and total oxygen consumed by each specimen. This would require measuring a specimen’s oxygen uptake before and after applying a photosynthetic inhibitor, but this was incompatible with our experimental design as the photosynthetic inhibition cannot be undone and we wanted to measure oxygen uptake rates twice, once before and once after the critical thermal maxima assessment. Note also that measuring specimens in the dark is not a viable way to measure a lack of photosynthetically-produced oxygen because these slugs have a circadian rhythm like other gastropods (Sandison 1967; Sokolova and Pörtner 2001) and exposure to darkness induces differences in their metabolic activity (Shirley and Findley 1978, Frankenbach *et al*. in review, EMJL unpublished results).

To measure oxygen uptake, each specimen was first placed in a respiratory chamber filled with 100% O_2_ saturated seawater at the same temperature to which it was exposed during the acclimation phase (detailed above). One chamber was filled with aerated seawater only to control for background microbial O_2_ consumption. The chambers were closed to prevent water exchange and placed in a water bath maintained at one of the four experimental temperatures (17°C or 22°C for measurements taken before the critical thermal limits were assessed or 27°C and 29°C for measurements taken afterward).

Different temperatures were used for the oxygen uptake measurements before and after the critical thermal maxima were assessed because we wanted to measure the oxygen uptake directly following the critical thermal maxima trials. However, maintaining slugs at their critical thermal maximum temperatures is lethal, so lowering the temperatures and allowing slugs to recover is crucial. Decreasing the temperature all the way down to their initial exposure temperatures (17°C and 22°C) would have taken multiple hours and we were concerned this delay would result in a measurement that missed the window when the animals were thermally stressed.

Each respiratory chamber was placed under the same light intensity to which the specimens were initially exposed to maintain the photoacclimation status of that specimen’s chloroplasts (100µmol m^-2^ s^-1^ or 200µmol m^-2^ s^-1^) and ensure that the slugs did not experience light stress during the experiment. Because we measured rates of oxygen uptake when exposed to light, each measurement comprises the net demand for oxygen (i.e. total oxygen demand minus photosynthetically-produced oxygen). The decrease in O_2_ saturation, i.e. the amount of O_2_ that was extracted from the water by the slug, was measured every 10 minutes for 1 hour, or until the oxygen saturation dipped below 70%, to prevent inadvertently measuring oxygen uptake under hypoxic conditions. The time of the experiment was noted for each specimen. Prior to each measurement, the light was briefly turned off to avoid any interference with the optical sensor. Directly before each measurement, each vial was gently inverted to mix the water and prevent any oxygen gradients from developing. Each respiratory chamber was sterilized with ethanol and thoroughly rinsed before and after each use. At the end of the experiment, the wet mass of each slug was measured by placing it on a sheet of aluminum foil. Excess water was removed with a tissue before each slug was transferred into a pre-massed petri dish filled with seawater on a scale.

To calculate the amount of seawater (and therefore oxygen) in each respiratory chamber, the volume of each slug was computed and then subtracted from the total volume of each chamber. Respiratory chambers of different sizes were used (2ml or 22ml), depending on the size of the slug. Since directly and accurately measuring the volume of each *E. viridis* specimen via displacement was impossible due to their small size, we used their wet mass to estimate their volume. Rather than assuming that their density equaled 1g/ml, we measured the volume and wet mass of the larger congeneric species, (*Elysia crispata* Mörch, 1863, n=10) to calculate its average density (1.25g/ml), which we used to convert wet mass into volume for *E. viridis* and then subtracted from the total volume of the respiratory chamber. Then, the initial and final oxygen saturation measurements (%) were converted to oxygen concentrations in mgO_2_/L seawater for each experimental temperature using an online calculator (https://water.usgs.gov/cgi-bin/dotables - U.S. Geological Survey, 2018). These values were then used to calculate the rates of oxygen uptake before and after the critical thermal maxima trials in mg O_2_ taken up by each individual per hour.

### Experimental phase - measuring critical thermal maxima (CT_first_ and CT_last_)

The protocol for measuring critical thermal maxima was adapted from Armstrong *et al*. (2019) and applied to every slug in every treatment for both Experiments 1 and 2. Each slug (n=8 intended per treatment) was placed in an individual bottle containing aerated seawater. These bottles were placed in a circulating water bath to control temperature. The same light intensity to which they had been exposed during the acclimation phase (100µmol m^-2^ s^-1^ or 200µmol m^-2^ s^-1^) was also provided. The temperature was increased by 1°C every 15 minutes (i.e. 0.0667 °C min^-1^) and slugs were observed continuously during this thermal ramping.

Preliminary trials were conducted to define two stages of heat stress in sea slugs, since definitions of the critical thermal maximum differ in the literature. In the first stage, starting from the initial exposure temperature (17°C or 22°C), the slugs were observed as they tried to escape by climbing up the walls of the bottle as the temperature increased. The end of the first stage was defined as the temperature at which a slug could no longer attempt to flee by climbing the walls of the test chamber, indicating it had lost voluntary muscle control. The temperature at this point is defined here as CT_first_. We also recorded a second stage of heat stress, where slugs lost the ability to contract muscles in response to external stimuli, reflecting neuronal dysfunction. This point was most reliably determined by gently prodding each slug’s rhinophores with a probe every few minutes. The temperature at which they could no longer contract a rhinophore when stimulated, i.e. they lost involuntary muscle control and nervous system function, was termed CT_last_. This point most closely corresponds to a heat-induced coma described by Lutterschmidt and Hutchison (1997).

Immediately upon reaching CT_last_, slugs were removed from the water bath and allowed to cool down by 1°C every 10 minutes to a temperature that facilitated recovery. Slugs that were initially exposed to 17°C during the acclimatory period were cooled down to 27°C for recovery and slugs initially exposed to 22°C during the acclimatory period were cooled down to 29°C for recovery. These temperatures were based on the average temperature at which all of the specimens recovered and none died of hyperthermia during a set of preliminary investigations. Heat tolerance plasticity was calculated using Acclimation Response Ratios (ARRs), which express how much heat tolerance is gained per degree of warm acclimation. ARRs were calculated as the increase in CT for every degree of warm acclimation, using the formula: ARR = (CT at 22°C - CT at 17°C)/5, as described by Gunderson & Stillman (2015).

### Experimental phase - measuring color and photosynthetic efficiency

The color of each specimen was recorded before its oxygen uptake and thermal maxima were recorded. *Elysia viridis* get their dark green coloration from their kleptoplasts and they fade to a terracotta color as kleptoplasts are digested during starvation. Four noticeable color stages were observed (dark green, pale green, pale brown, terracotta).

Five specimens of each color were dark-acclimated for 15 minutes before being measured with a Pulse Amplitude Modulated (PAM) fluorometer (Mini-PAM, Walz GmBH, Germany), to determine the average photosynthetic efficiency (Fv/Fm) at each color stage and to assess if specimens had ingested chloroplasts that could photosynthesize and the photosynthetic efficiency of these chloroplasts, as has been observed in previous studies (see e.g. Vieira *et al*., 2009; Serôdio *et al*., 2014; Laetz and Wägele., 2018).

### Statistical analyses

All analyses were conducted in R-studio (RStudio Team, 2020) based on R version 3.6.1 (R Core Team 2019) using the packages *matrixstats* (Bengtsson, 2017), *lme4* (Bates et al. 2015), *MuMin* (Bartoń, 2022), *lmerTest* (Kuznetsova et al. 2017), *ggplot2* (Wickham, 2016), *dplyr* (Wickham et al. 2022), and *visreg* (Breheny & Burchett, 2017).

The R-script we generated and all datasheets have been uploaded to DataverseNL (currently a temporary link, final link will be provided if the manuscript is accepted: https://dataverse.nl/privateurl.xhtml?token=56681d13-8b25-45ba-80c4-4ad619eaf2a3). We used general linear models to investigate how our treatments affected each slug’s critical thermal maxima. In both experiments, the two stages of heat stress (CT_first_ and CT_last_) were positively correlated (Experiment 1: B=0.221; t_1,111_ = 4.66; P < 0.001 (Fig. S1A) and Experiment 2: B=0.234; t_1,61_ = 3.468; P < 0.001 (Fig. S1B), where B is the slope, t is the t-value and its subscripts are the degrees of freedom for the numerator and denominator separated by a comma, and P is the probability value). Since they were correlated in both experiments, we merged data for CT_first_ and CT_last_, adding a binary dummy variable (CT_type) to distinguish between them for each experiment prior to analysis.

In Experiment 1, we included the temperature to which slugs were exposed, algal food source, and starvation period as fixed effects (Fig. S2A). We also included the body mass of each individual slug, as previous studies have demonstrated that heat tolerance may vary with body size in aquatic ectotherms (Leiva *et al*., 2019). A purely additive model already explained 83.6% (adjusted R ^2^ = 0.836) of the variation in critical thermal maxima, but we also tested whether interactions between predictor values improved the model. To test whether behavioral responses to heat stress (indicated by CT_first_) and neuronal dysfunction (indicated by CT_last_) were controlled by different drivers, we considered interactions between our binary dummy variable and the other fixed factors. Additionally, we considered interactions between experimental conditions, notably starvation duration and algal source to test whether slugs fed with different algae responded differently to starvation. To account for non-linear effects, we also included a quadratic term for starvation duration. To prevent overfitting, we considered only 2-way interactions and simplified the model by excluding non-significant interactions and interactions that contributed little to the explained variation. This resulted in a final model that explained 85.5% (adjusted R ^2^ = 0.855) of the variation (Table S1).

In Experiment 2, we included starvation period and light intensity as fixed effects. A full model allowing interactions explained 82.2% (adjusted R ^2^ = 0.822) of the variation in critical thermal maxima. We simplified this model by removing non-significant interactions resulting in a model that explained 81.8% (adjusted R ^2^ = 0.818) of the variation. A comparison of these models with AIC indicated that the simpler model is a better fit so this is the model on which our results are based (Table S2).

When analyzing rates of oxygen uptake for both Experiments 1 and 2, we first accounted for effects of body mass. The body mass of our slugs varied across two orders of magnitude. Both initial differences in size resulted from the algal food source they consumed during the acclimation period (Fig. S3), and decreases in body mass during starvation contributed to the observed variation in body mass. Since rates of oxygen uptake scale allometrically with body mass (White *et al*., 2007; Rubalcaba *et al*., 2020), we could not express rates of oxygen uptake on a per gram basis. Instead, we corrected rates of oxygen uptake using an empirically derived scaling exponent. Because body mass was statistically related to two treatment conditions (the initial temperature to which they were exposed (termed acclimation temperature in these analyses) and starvation duration), we needed to ensure that our empirically derived scaling exponent did not reflect variation in body mass across these treatment conditions. Therefore, we calculated the exponent by fitting a random intercept for each combination of acclimation temperature and starvation period (i.e. = 2 temperatures x 5 starvation intervals = 10 levels), thus focusing on the effect of within-treatment variation in body mass. This yielded a scaling exponent of 0.792+0.05 S.D. (Fig. 3A), which agrees well with known values for ectotherms in the literature (White *et al*., 2007; Verberk *et al*., 2020). Effects of body mass explained ∼55% of the variation in absolute oxygen uptake rates.

**Figure 3.**
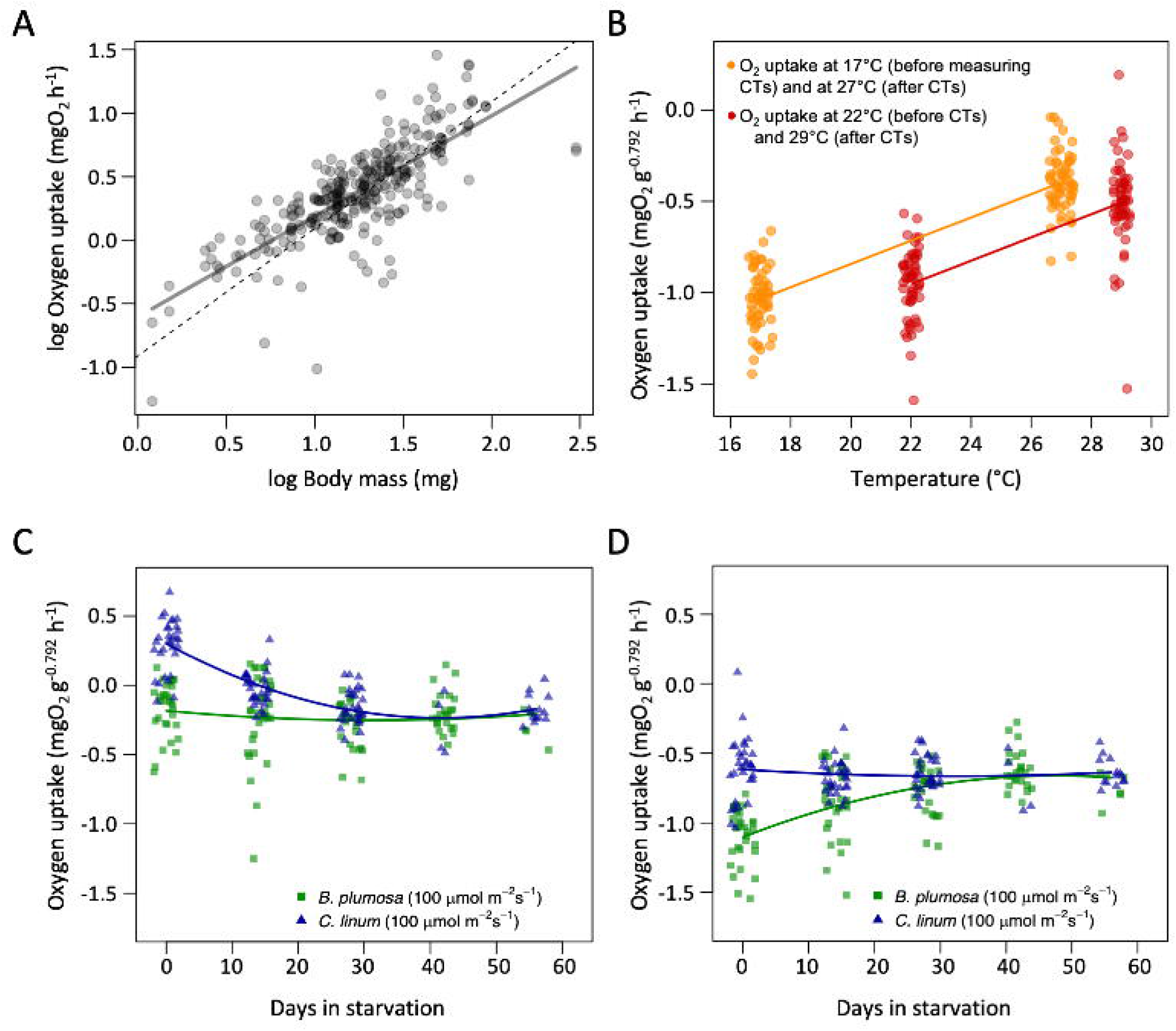
Mass specific rates of oxygen uptake. A) The allometric scaling relationship between log body mass (mg) and log oxygen uptake (mgO_2_ h^-1^). The solid line indicates the empirically derived scaling exponent of 0.792+0.05 S.D., while the dotted line indicates isometry, i.e. a scaling exponent of 1. This scaling exponent was used to calculate the mass-specific rates of oxygen uptake rates in Experiments 1 and 2. B) Variation in mass specific rates of oxygen uptake at each testing temperature for Experiment 1. Specimens exposed to 17°C (orange) were measured before exposure to acute heat stress (critical thermal maxima trials) and after exposure to acute heat stress/recovery at 27°C. Specimens exposed to 22°C (red) were measured before exposure to acute heat stress (critical thermal maxima trials) and after exposure to acute heat stress/recovery at 29°C. C) The amount of oxygen taken up during starvation before the critical thermal maxima trials and D) after the critical thermal maxima trials for specimens in Experiment 1 (fed *B. plumosa* (dark green squares) and *C. cf. linum* (dark blue triangles)). Note that individual data points are slightly jittered on the x-axis. These partial residual plots illustrate the relationship between the response variable and a given independent variable while accounting for the effects of other independent variables in the model (Table S3). The lines are regression lines and the shaded bands are 95% confidence intervals.

Next, we analyzed these mass-corrected rates of oxygen uptake (mg O_2_ g^-0.792^ h^-1^), using linear mixed effects models for both Experiments 1 and 2. For each individual slug we measured oxygen uptake rates twice, before and after exposure to acute thermal stress where critical thermal maxima were assessed (Fig. 2; detailed in the subsection: “*Experimental phase - calculating the rate of oxygen uptake* ”. Both measurements were included in a single model, adding “HS” (heat shock) as a binary dummy variable to distinguish between oxygen uptakes measurements made pre- and post-the critical thermal maxima trials. This approach allowed us to better disentangle the effects of the initial temperature to which they were exposed and the temperature at which they were tested because different test temperatures were employed before (17°C and 22°C) and after (27°C and 29°C) their critical thermal maxima were measured. This was accounted for by correcting for the test temperature via the coefficient.

In Experiment 1, we started with a simple additive model that included the fixed effects: acclimation temperature (17°C or 22°C), the test temperature (17°C, 22°C, 27°C, or 29°C), a quadratic term for starvation duration to account for non-linear effects, the algae food source and our dummy variable (HS) indicating if the measurement for oxygen uptake was taken before or after critical thermal maxima were measured. We also included individual identity as a random factor because each individual was measured twice (before and after their critical thermal maxima were measured). This additive model explained 27% of the variation (marginal R ^2^ = 0.22, conditional R ^2^ = 0.27). To try to improve this model, we included an interaction between starvation period and algal source to test whether slugs fed with different algae responded differently to starvation, which we predicted due to the different amounts of time *E. viridis* can retain chloroplasts from these algal genera. We also tested whether responses to algal source and starvation differed for oxygen uptake rates measured before or after the thermal tolerance assay (i.e. pre- and post-heat-shock). To do this, we compared a model that included interactions between heat-shock and both starvation and algal source against a model that did not include these interactions. The data best supported a model that explained 49% of the variation (marginal R ^2^ = 0.42, conditional R ^2^= 0.49) and is reported in our results (Table S3).

In Experiment 2, we built a model with the following fixed effects: light intensity, starvation days (again as a polynomial), and the dummy variable (HS) to distinguish between measurements of oxygen uptake before and after critical thermal maxima were measured. We also included slug identity as a random factor since each slug was measured twice (before and after its critical thermal maxima were measured). Since the slugs in this experiment were only fed *B. plumosa* and exposed to 17°C, algal food source and acclimation temperature were not included in this analysis. In addition, rather than including test temperature (17°C or 27°C), rates of oxygen uptake were standardized to an average temperature of 22°C, using the thermal sensitivity derived from Experiment 1. This solved the problem that the effect of test temperature and heat shock were collinear in Experiment 2 (slugs were measured at 17°C before the heat-shock and at 27°C after the heat shock). This model already explained 64% of the variation (adjusted R ^2^) and had an AIC value of -29.40. We then simplified this model by removing non-significant interactions, producing a simpler model that also explained 64% of the variation and had a lower AIC value (-32.55). This simpler model is reported in our results and contains light intensity, HS, and starvation days as fixed effects with an interaction between HS and starvation days (Table S4).

## Results

### Specimen mass, color and photosynthetic activity during starvation

All slugs had a dark green color due to the chloroplasts they had previously incorporated when they entered the lab and began the acclimation phase. They remained dark green throughout the acclimation phase during which they were allowed to feed *ad libitum*, but once they entered the starvation period, color changes were observed. After an extended period of starvation, many of the slugs had terracotta coloration, indicating they had lost most of their chloroplasts (Fig. 4A). Parallel decreases in photosynthetic efficiency were observed during starvation, as indicated by decreasing Fv/Fm (PAM fluorometry) measurements (Fig. 4B). Slugs also decreased in length and mass during the starvation period (Fig. 4B). In Experiment 1, slugs fed *C.* cf. *linum* were larger than slugs fed only *B. plumosa* after the acclimation phase (Fig. S3), however these slugs also lost mass faster than slugs fed *B. plumosa* during the starvation period. For example, at 28 days in starvation, slugs fed *C.* cf*. linum* averaged 18.91 ± 8.08 mg while those fed *B. plumosa* averaged 27.29 ± 14.56 mg (at 17°C and 100µmol m^-2^ s^-1^).

**Figure 4.**
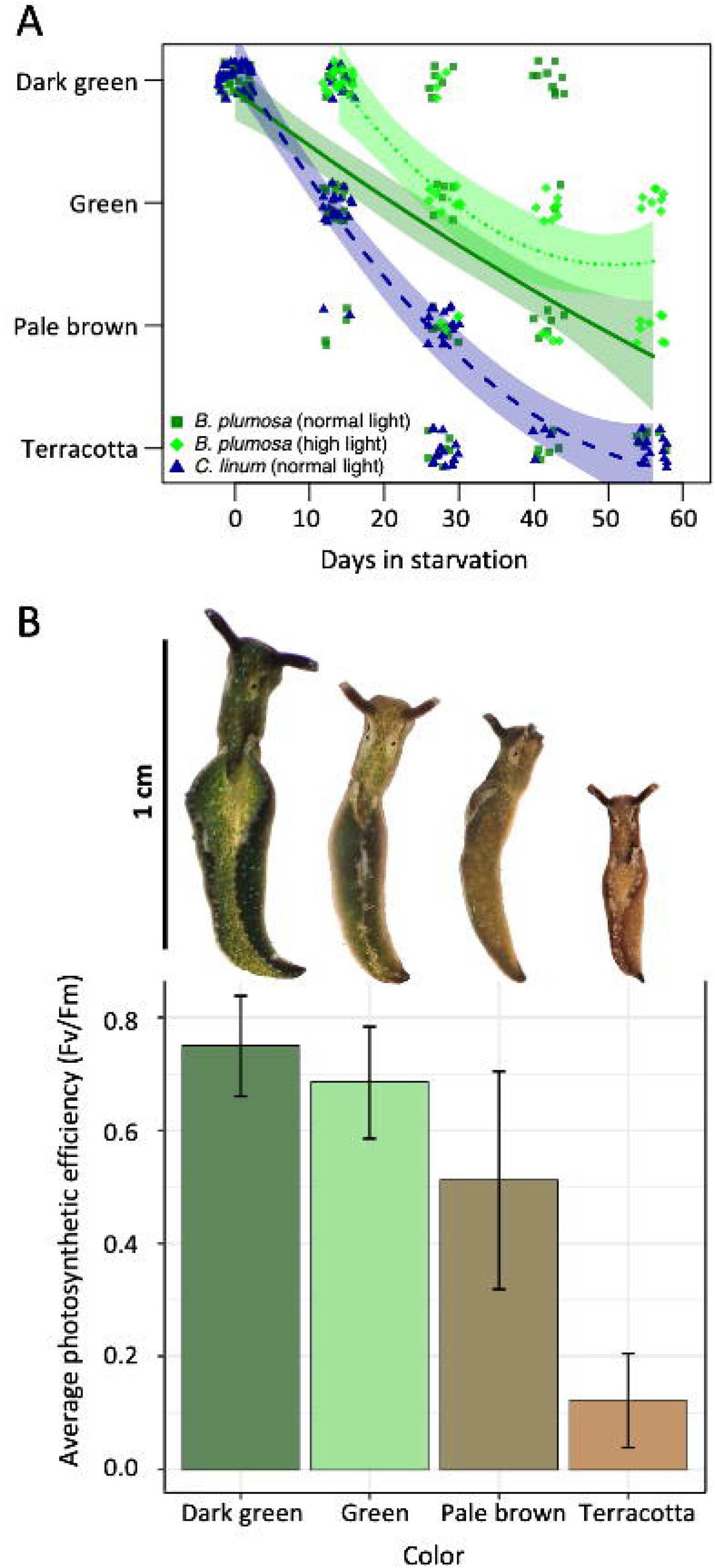
Change in color during starvation and the Fv/Fm at each color stage for specimens exposed to 17°C. A) Loss of chloroplasts during starvation causes a color change from dark green to green to pale brown and finally terracotta. The rate of bleaching and hence loss of chloroplasts differed between the two algal food sources at 17°C. Specimens fed *B. plumosa* under normal light are depicted in dark green squares, specimens fed *B. plumosa* under high light are shown in light green diamonds and specimens fed *C. cf. linum* are illustrated in dark blue triangles. Data points are jittered on both axes to increase visibility. Model predictions are from a model with starvation period as a quadratic function. The lines are regression lines and the shaded bands are 95% confidence intervals. B) An example of a slug depicting each color and the average Fv/Fm value for each color category. Each column height depicts the mean Fv/Fm (n=5) and the lines represent the standard deviation for each column.

### Heat tolerance

Experiment 1 examined how higher temperature, starvation and algal food source impact a slug’s critical thermal maxima. A model that included the effects of acclimation temperature, CT type (CT_first_ or CT_last_), body mass, algal food source, and starvation duration, including several interactions, had an adjusted R ^2^ of 0.86 (Table S1). In addition, variation in critical thermal maxima could be related to effects of starvation, algae, light and initial exposure temperature (Fig. 5; Table S1). Slugs exposed to 22°C (the higher temperature) showed improved heat tolerance compared to slugs exposed to 17°C (Fig. 5A), and this significantly differed between CT_first_ and CT_last_, (t_1,213_ = 3.424; P<0.001) and varied with body mass (Fig. 5B; t_1,213_ = 2.718; P = 0.0071). For a slug of average size, the Acclimation Response Ratios (ARR, which expresses how much heat tolerance (in °C) is gained for every degree of warm acclimation) was higher for CT_first_ (ARR =0.225), than for CT_last_ (ARR =0.036). This difference in heat tolerance with acclimation temperature was most pronounced in larger slugs, giving rise to a negative relationship with body mass, such that larger-bodied slugs exhibited reduced heat tolerance, but only for slugs exposed to 17°C (Fig. 5B). Starvation affected critical thermal maxima depending on the algae the slug had consumed and on the temperature to which slugs were initially exposed (17°C or 22°C). At the start and at the end of the starvation period, slugs exhibited improved heat tolerance when fed *C.* cf*. linum* (Fig 5C; Table S1), whereas an improved heat tolerance due to warm acclimation was observed mainly in the middle of the starvation period (Fig. 5D).

**Figure 5.**
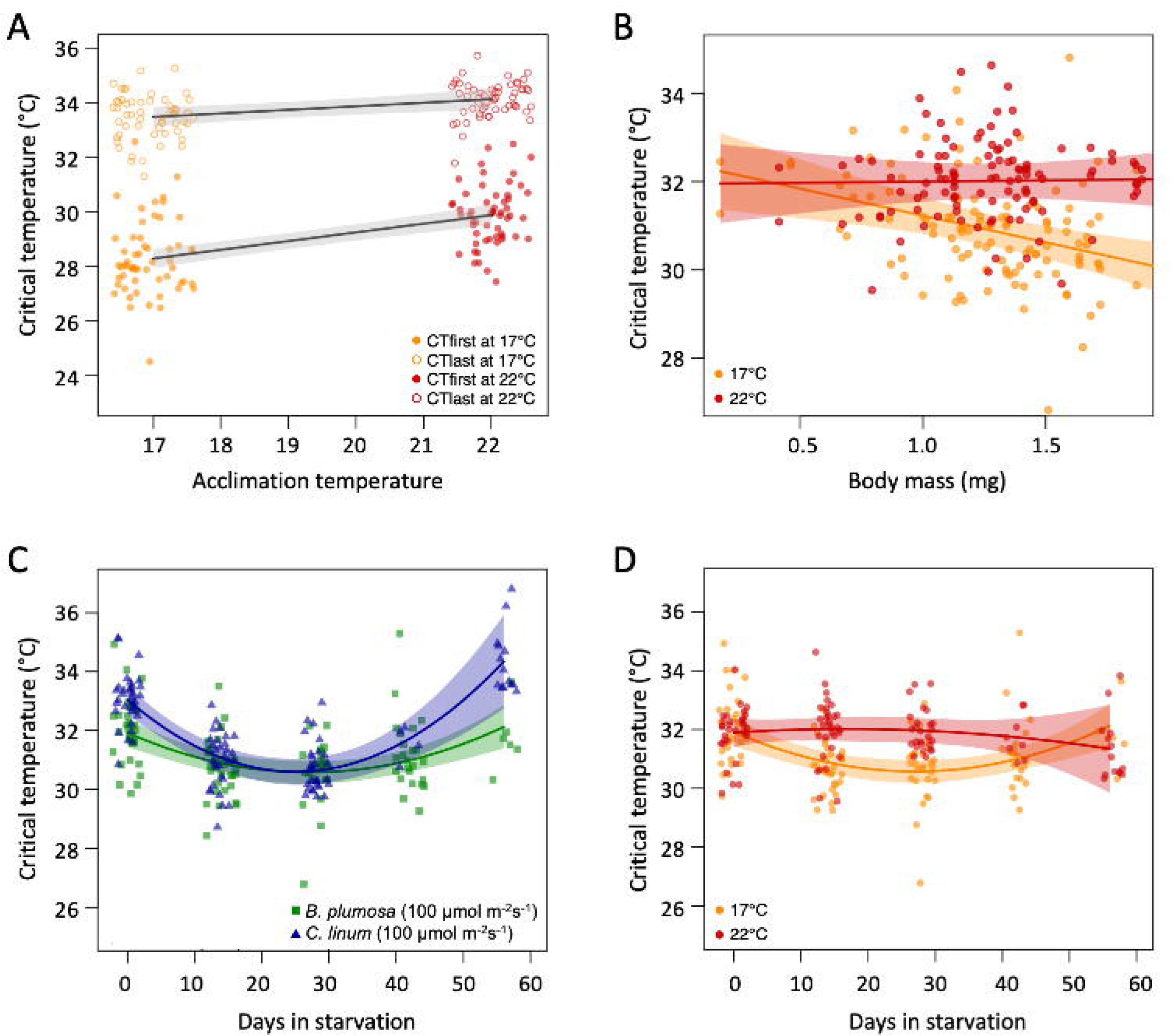
Critical thermal maxima in E. viridis during Experiment 1. A) Variation in heat tolerance (CT_first_ (closed circles) and CT_last_ (open circles)) differed due to initial exposure temperature (17°C in orange, 22°C in red) and B) log-transformed body mass (mg). C) Thermal tolerance also varied due to starvation duration, food and light conditions. *B. plumosa* fed slugs under normal light are shown via dark green squares, and *C.* cf. *linum* fed slugs under normal light are represented by dark blue triangles. D) Thermal tolerance for each of the experimental temperatures during starvation. Note that individual data points are slightly jittered along the x-axis for (A) and (C). These partial residual plots illustrate the relationship between the response variable and a given independent variable while accounting for the effects of other independent variables in the model (Table S2). The lines are regression lines and the shaded bands are 95% confidence intervals.

Experiment 2 examined the effect of light intensity on critical thermal maxima in slugs exposed to 17°C and fed *B. plumosa.* There was a significant interaction between light intensity and starvation period (Table S2). Under 200µmol m ^-2^ s^-1^ (the higher light intensity), they initially exhibited a better heat tolerance on average than those at 100µmol m^-2^ s^-1^, but the effect of light intensity disappeared towards the end of the starvation period (Fig. 6A).

**Figure 6.**
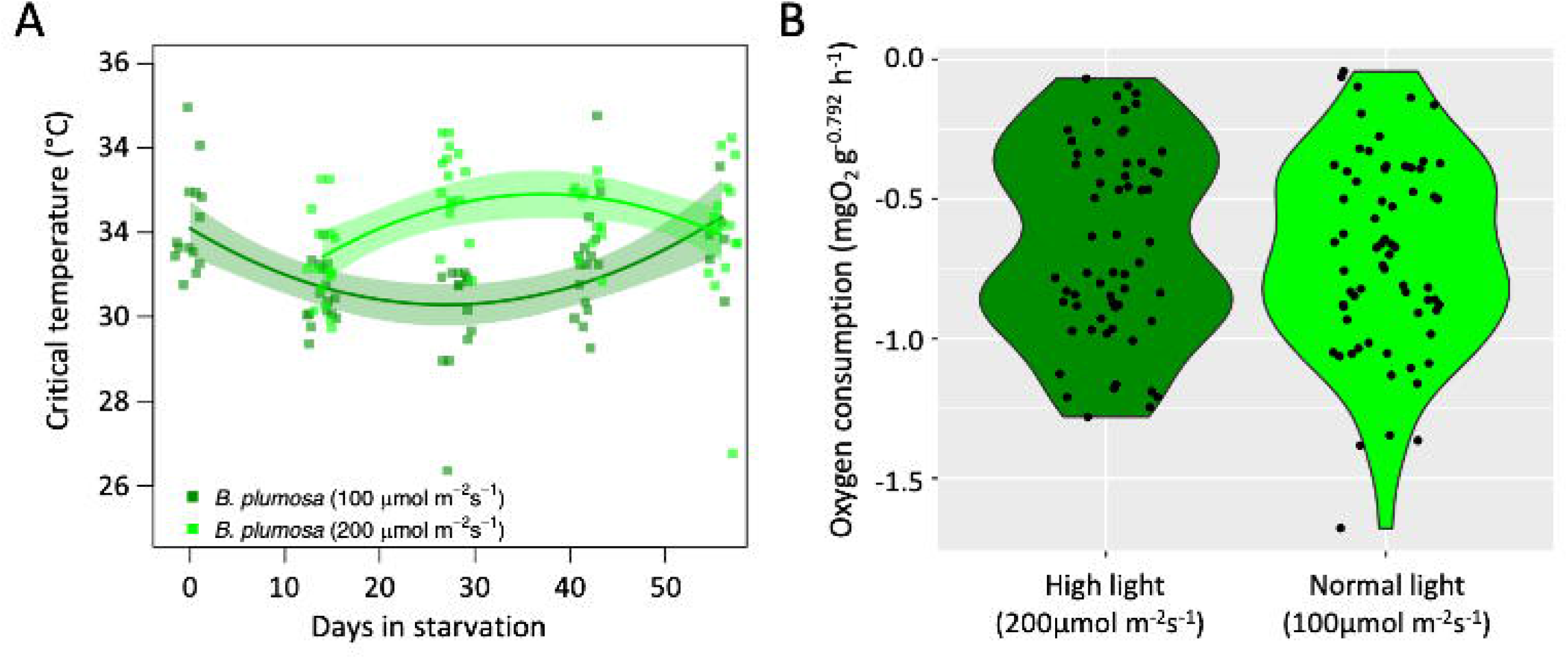
Critical thermal maxima and oxygen uptake under different light intensities for E. viridis kept at 17°C and fed B. plumosa (Experiment 2). A) Critical thermal limits in slugs exposed to 200µmol m^-2^ s^-1^ and 100µmol m^-2^ s^-1^ during starvation. Slugs exposed to 200µmol m^-2^ s^-1^ initially exhibited a better heat tolerance on average than those at 100µmol m^-2^ s^-1^, but this disappeared towards the end of the starvation period. B) Oxygen uptake in slugs exposed to 200µmol m^-2^ s^-1^ and 100µmol m^-2^ s^-1^ did not differ. Note that individual data points are slightly jittered along the x-axis in (A) and (B).

### Rate of oxygen uptake

The oxygen concentration decreased in every respirometry chamber throughout the experimental period, despite ongoing photosynthetic activity at the beginning of the starvation period, indicating that oxygen uptake via aerobic respiration outpaces oxygen production via photosynthetic activity. A strong allometric scaling relationship was evident between rates of oxygen uptake and body mass (Fig 3A), so our subsequent analyses focused on mass-corrected rates of oxygen uptake (mg O_2_ g^-0.792^ h^-1^; see Methods). The best model for Experiment 1 included acclimation temperature, test temperature, HS (the dummy variable we used to distinguish MO2 measurements taken before or after the critical thermal maxima trial), algal food source and starvation duration as fixed effects and included several interactions. Rates of oxygen uptake were reduced in slugs exposed to a warmer temperature (t =-3.309; P = 0.001) and increased with measurement temperature (Fig 3B; t =3.476; P < 0.001). Slugs fed with *C. linum* exhibited higher rates of oxygen uptake than those fed with *B. plumosa* at the start of the starvation period, but as starvation progressed, oxygen uptake rates declined to the same level (Fig. 3C; Table S3). Oxygen uptake rates were lower after the critical thermal maxima were measured, i.e. the slugs took up less oxygen after exposure to acute heat shock (t_1,109_=-4.036; P<0.001; Fig. 3D). This reduction was most pronounced at the start of the starvation period (note that the effect of a higher measurement temperature post heat-shock is accounted for in Fig. 3D).

Experiment 2 examined the effects of light intensity on oxygen uptake in slugs kept at 17°C and fed *B. plumosa*. Differences in light intensity did not affect oxygen uptake rates (Fig. 6B; Table S4).

## Discussion

### Heat tolerance and oxygen

*Elysia viridis* heat tolerance improved following exposure to 22°C, relative to slugs exposed to 17°C. This acclimatory ability to improve heat tolerance is widely investigated in ectotherms (Gunderson and Stillman, 2015), including sea slugs (Armstrong *et al*., 2019). Our results show that the ARR varied between 0.036 and 0.225, falling within the range of values previously reported for sea slugs (Armstrong *et al*., 2019). Perfect acclimation (i.e an ARR value of 1) would mean that the animal can keep increasing its heat tolerance in tandem with warming of its habitat. Since acclimation was not perfect (ARR < 1), there appears to be a limit in how plasticity in heat tolerance can help buffer *E. viridis* against the highly variable temperature fluctuations that they experience in their uppermost subtidal habitats, although the exact limit may be time-dependent (Semsar-Kazerouni and Verberk, 2018). Armstrong *et al*. (2019) compared the acclimatory ability to improve heat tolerance by calculating ARRs and using a critical thermal limit that aligns with our definition of CT_first_ in nudibranch sea slugs (Gastropoda: Heterobranchia). They found that the higher tolerance limits exhibited by heat-tolerant species were less plastic than the lower tolerance limits exhibited by heat-sensitive species. Here we observe a similar pattern within a species: acclimation temperature had a stronger effect on CT_first_, an endpoint associated with lower temperatures that reflects loss of voluntary muscle control, while CT_last_, an endpoint associated with higher temperatures, and reflects neuronal dysfunction, was less responsive to acclimation temperature (Fig. 5A). One way to explain this is that with increasing intensity of heat stress, more and more physiological mechanisms (e.g. oxygen provisioning, protein structure, membrane function) will become prone to failure and physiological adjustment of all these mechanisms will become increasingly difficult. Thus, earlier critical points, which are recorded at lower temperatures, likely involve fewer mechanisms which makes them more malleable in terms of physiological acclimation.

Several observations are consistent with the hypothesis that oxygen becomes limiting when animals reach their thermal maxima, necessitating a switch to anaerobic metabolism. Oxygen availability in seawater has a demonstrable effect on heat tolerance in gastropods (Koopman *et al*., 2016; Hoefnagel and Verberk, 2017). While most marine gastropods have a gill or gills, most sacoglossans including *E. viridis* take up oxygen through their epidermis (Clark *et al*., 1981; Graham, 1990; Neusser *et al*., 2019), a mode of respiration that is especially prone to suffer from oxygen limitation at thermal extremes (Verberk and Bilton, 2013). Moreover, smaller individuals have higher surface area to volume ratios, and we observed that they are more heat tolerant than larger individuals, at least when exposed to 17°C (Fig 5B). A previous meta-analysis also showed that larger aquatic ectotherms exhibited lower heat tolerance, especially in longer ramping trials where the efficacy of anaerobic metabolism is reduced (Leiva *et al*., 2019). In this study, we increased the temperature by 1°C every 15 minutes so the ramping trials took multiple hours. We also observed an increase in heat tolerance for individuals exposed to high-light conditions (∼2°C higher than slugs in low-light conditions), which could be due to increased oxygen production from photosynthesis. Widespread distribution of this oxygen throughout the animal in the highly branched digestive gland could alleviate oxygen limitation. An oxygen interpretation is also consistent with our observation that critical thermal maxima were comparable across light levels at the very end of the starvation period, when photosynthesis had ceased due to chloroplast degradation and all specimens were bleached.

### Food and light intensity

In their natural habitats, *E. viridis* is often the predominant macroscopic herbivore and specimens can be found in great abundance near or on their food algae (Hinde and Smith, 1975; Trowbridge *et al*., 2008). This study demonstrates that the Dutch population of *E. viridis* from Bommenede naturally feeds on *B. plumosa* and that it can retain chloroplasts from *B. plumosa* for more than a month, which expands the list of algal species whose chloroplasts can be retained by these kleptoplastic slugs. The abundance and availability of algal species on which these slugs feed can vary greatly throughout the year in the North Sea (Baumgartner and Toth, 2014, EMJL unpublished observation). Data on year-round abundance of *B. plumosa* in Zeeland is however lacking, and algal species can be unavailable periodically if conditions for germination are unfavorable (Rietema, 1969, 1970). Food availability may therefore be a limiting factor for temperate, stenophagous species like *E. viridis* and this could have selected for the evolution of kleptoplasty or the ability to feed on multiple algal food sources (Wägele and Martin, 2014).

Almost every specimen exposed to 22°C had a bleached appearance due to chlorophyll breakdown by 28 days in starvation, whereas many specimens exposed to 17°C retained bright green coloration from chlorophyll for more than 42 days. Since kleptoplasts provide some energy to starving *E. viridis* via photosynthesis and this energy may help them withstand periods of food unavailability (Cartaxana *et al*., 2019), the premature breakdown of chlorophyll earlier in the starvation period at 22°C could have serious energetic consequences for starving slugs during times of food unavailability in a warmer world.

Our results demonstrate that *E. viridis* fed exclusively *C.* cf. *linum* could retain functional chloroplasts for more than 17 days in starvation, contrasting reports that hypothesized *E. viridis* could not retain *C.* cf*. linum* chloroplasts (Clark *et al*., 1990; Trowbridge and Todd, 2001). Moreover, slugs fed exclusively on *C.* cf*. linum* grew almost twice as long and they were ∼3 times heavier than members of the same population that were maintained on *B. plumosa*. Baumgartner *et al*. (2014) found that *E. viridis* gained more mass when fed *C. melagonium*, aligning with our observations and indicating that *Chaetomorpha* spp. are particularly nutritious for *E. viridis*, even if certain populations rarely consume this alga in nature (Trowbridge *et al*., 2008). Other sacoglossan species have also been reported to thrive on food algae on which they are seldom observed eating in nature (Clark, 1975; Barber *et al*., 2021). Before their critical thermal maxima were measured, slugs fed with *C.* cf*. linum* exhibited higher rates of (mass-corrected) oxygen uptake than those fed with *B. plumosa*, possibly due to the increased growth and locomotion we observed in these *C.* cf. *linum* fed specimens. These differences disappeared with longer starvation duration, probably due to the faster rate of chloroplast breakdown in specimens fed *C.* cf*. linum*. Despite their large initial size, *E. viridis* specimens fed *C.* cf*. linum* lost both mass and photosynthetic capacity faster than specimens fed *B. plumosa*, indicating that kleptoplasts from *C.* cf*. linum* are less advantageous in terms of nutrition and oxygen production during extended starvation. In contrast, specimens fed *B. plumosa* and exposed to normal light retained functional chloroplasts the longest and exhibited the lowest weight loss during starvation.

### Oxygen uptake

Slug metabolism was strongly related to body mass according to a power law with a mass scaling exponent of 0.79, which agrees with empirical estimates for ectotherms (White *et al*., 2007; Verberk *et al*., 2020). Since the effect of body mass was allometric (i.e. a scaling exponent <1), mass specific oxygen uptake (i.e. dividing respiration rates by mass) would underestimate metabolism in larger individuals. Therefore, we adjusted oxygen uptake rates using the empirical mass scaling exponent. This corrected for indirect effects of temperature, food and starvation on body size and instead isolated direct effects. Slugs acutely exposed to warmer temperatures increased their oxygen uptake rates, but after exposure to warmer conditions, oxygen uptake rates were downregulated (Fig. 3B) to compensate for this thermodynamic effect (Seebacher *et al*., 2015). In every trial, we measured net oxygen uptake, indicating that photosynthesis in *E. viridis* did not produce enough oxygen to support aerobic respiration. It was not possible to determine how much oxygen was produced via photosynthesis in each specimen, since repeat measures of each animal in the dark or with when the animals were subjected to photochemical inhibition would have required repeat measures or doubling the sample size, both of which were not feasible during this study. This means that the “true” amount of oxygen needed to support aerobic respiration will require further study. Previous work by Dionísio *et al*., (2018) on this species and its tropical congener *E. crispata* showed net oxygen production via photosynthesis which contrasts our results. However, it is not clear on which algal species this population of *E. viridis* specimens was feeding since they were provided both *B. plumosa* and *Codium tomentosum* Stackhouse 1797, meaning this discrepancy could be due to the incorporation of chloroplasts from *C. tomentosum*.

During starvation, *E. viridis* digest their incorporated chloroplasts (Laetz *et al*., 2016; Rauch *et al*., 2018) so the sampling points that occurred late in the starvation period (42 and 56 days) surveyed bleached animals, which were almost certainly chloroplast-free or only contained degraded chloroplast remnants. This assertion is corroborated by the extremely low or unmeasurable photosystem II activity we measured with PAM fluorometry. This indicates that oxygen production due to photosynthetic activity has also ceased late in starvation. We expected to observe increased oxygen uptake rates as starvation progressed as slugs take up more oxygen from the seawater to compensate for the declining oxygen availability from photosynthesis, however, oxygen uptake rates decreased during starvation. When facing starvation, animals may conserve ATP by metabolic suppression (de Vries *et al*., 2015; Semsar-Kazerouni *et al*., 2020), which likely explains the abrupt decline in oxygen uptake we observed as starvation began (Fig. 3C). Metabolic suppression is a survival strategy in which an organism suppresses non-critical cellular processes to reduce its oxygen uptake and conserve energy in an attempt to ensure long-term survival during acute environmental stress (Sokolova and Pörtner, 2001).

Locomotion, growth, and reproduction may all be curtailed to save energy. Metabolic suppression likely also explains the decrease in oxygen uptake observed when comparing oxygen uptake rates measurements before and after the heat tolerance assay (compare Fig. 3C and Fig. 3D). This strategy is often observed in intertidal gastropods that have shells in which to retreat during periods of thermal stress (Sokolova and Pörtner, 2001, 2003; Marshall *et al*., 2011), but examinations of shell-less gastropods (slugs) are limited.

### Conclusions

The data presented here indicate that these slugs will be able to cope with gradual and acute warming of their habitat. To deal with temperature increases, they can downregulate their metabolism and increase their heat tolerance via plasticity. All but three of the *E. viridis* specimens that experienced acute heat stress were able to recover from the loss of voluntary muscle control and neuronal dysfunction associated with heat shock, indicating that exposure to their thermal maxima is rarely lethal. This species lives in shallow coastal waters where exposure to heat and high light intensity will be linked, leading to internal oxygen production which may help alleviate some degree of oxygen limitation, even though photosynthetic oxygen production does not provide all of the oxygen needed to fuel aerobic respiration. Provided their algal food species remain plentiful and do not experience dramatic seasonality shifts due to warmer conditions so the slugs are not faced with excessive starvation periods, protracted warm periods should not be a problem for *E. viridis*. Finally, the surface temperature at our collection site varied seasonally between ∼5 and 23°C (Rijkswaterstaat, 2020). *Elysia viridis* live above the thermocline for most of the year, but migrated below it in July when the surface temperature exceeded 17°C. This avoidance behavior and the physiological adjustments in oxygen uptake and heat tolerance suggest that *E. viridis* with access to their food algae are capable of withstanding the warmer, deoxygenated waters in the future.

## Acknowledgements

We would like to thank Britas Klemens Eriksson for allowing us to use his lab facilities and culture rooms, Stella Bos for her help culturing algae, and Jan Veldsink for help ordering supplies, Maria van Leeuwe for lending us her PAM, Laura Govers and Gabriela Maldando for lending us a quantum sensor, and Hinke Tjoelker for managing administration (all University of Groningen).

We are grateful to the Dutch Research Council (NWO), who financed this work as part of the projects VI.Veni.202.218 (awarded to EMJL) and NWO-VIDI 016.161.321 (awarded to WCEPV).

## Declarations

The authors declare they have no competing interests.

## Ethical Care

All specimens were treated according to best practices, which aim to minimize animal suffering whenever possible. Following each trial, specimens were allowed to recover. When needed, animals were dispatched via flash freezing to minimize suffering.

## Data Availability

All of our data and R-scripts will be made publicly available in the online data repository, DataverseNL under a CC-BY license (the link below is a temporary https://dataverse.nl/privateurl.xhtml?token=56681d13-8b25-45ba-80c4-4ad619eaf2a3

## Supplementary Materials

**Table S1.** Summary table of the final model used in evaluating the effects of temperature, starvation and algal food source on critical thermal maxima (Experiment 1).

|  | Estimate | Std. Error | t value | Pr(>\|t\|) |
| --- | --- | --- | --- | --- |
| (Intercept) | 40.30825 | 2.48701 | 16.207 | **< 2e-16** |
| CT_typeVoluntary | -8.43269 | 1.08534 | -7.770 | **3.29e-13** |
| acclim_temp | -0.28943 | 0.12702 | -2.279 | **0.023677** |
| BM | -5.55302 | 1.84100 | -3.016 | **0.002870** |
| AlgaeCl | 0.65807 | 0.15995 | 4.114 | **5.55e-05** |
| poly(Starvation_days, 2)1 | -2.14047 | 11.63261 | -0.184 | 0.854184 |
| poly(Starvation_days, 2)2 | 44.66616 | 11.02956 | 4.050 | **7.18e-05** |
| CT_typeVoluntary:acclimtest_temp | 0.18923 | 0.05527 | 3.424 | **0.000740** |
| acclimtest_temp:BM | 0.25489 | 0.09377 | 2.718 | **0.007105** |
| acclimtest_temp:poly(Starvation_days, 2)1 | 0.01108 | 0.63228 | 0.018 | 0.986038 |
| acclimtest_temp:poly(Starvation_days, 2)2 | -2.12263 | 0.59996 | -3.538 | **0.000495** |
| AlgaeCl:poly(Starvation_days, 2)1 | 1.95868 | 3.29130 | 0.595 | 0.552403 |
| AlgaeCl:poly(Starvation_days, 2)2 | 9.54410 | 2.99593 | 3.186 | **0.001661** |

**Table S2:**
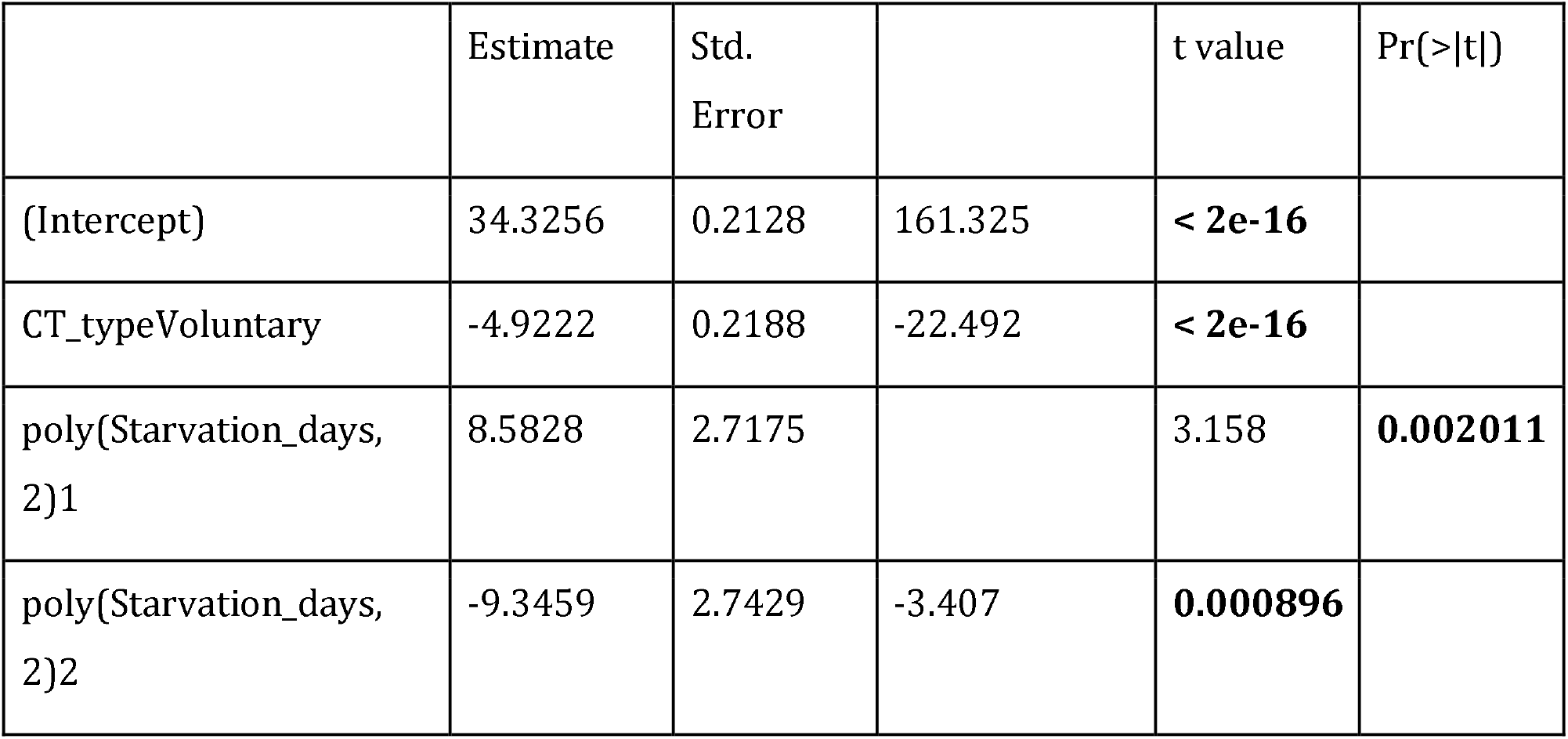
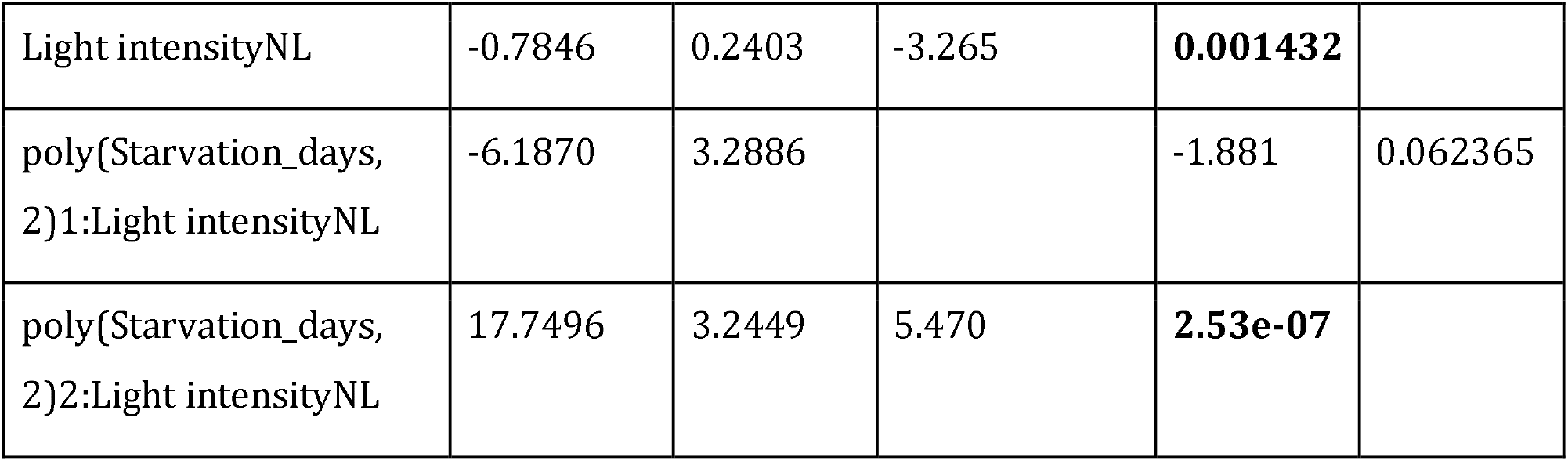
Summary table of the final model used in evaluating the effects of light intensity on critical thermal maxima for Experiment 2.

|  | Estimate | Std. Error | t value | Pr(>\|t\|) |
| --- | --- | --- | --- | --- |
| (Intercept) | 34.3256 | 0.2128 | 161.325 | **< 2e-16** |
| CT_typeVoluntary | -4.9222 | 0.2188 | -22.492 | **< 2e-16** |
| poly(Starvation_days, 2)1 | 8.5828 | 2.7175 | 3.158 | **0.002011** |
| poly(Starvation_days, 2)2 | -9.3459 | 2.7429 | -3.407 | **0.000896** |

|  |  |  |  |  |  |
| --- | --- | --- | --- | --- | --- |
| Light intensityNL | -0.7846 | 0.2403 | -3.265 | **0.001432** |  |
| poly(Starvation_days, 2)1:Light intensityNL | -6.1870 | 3.2886 |  | -1.881 | 0.062365 |
| poly(Starvation_days, 2)2:Light intensityNL | 17.7496 | 3.2449 | 5.470 | **2.53e-07** |  |

**Table S3.**
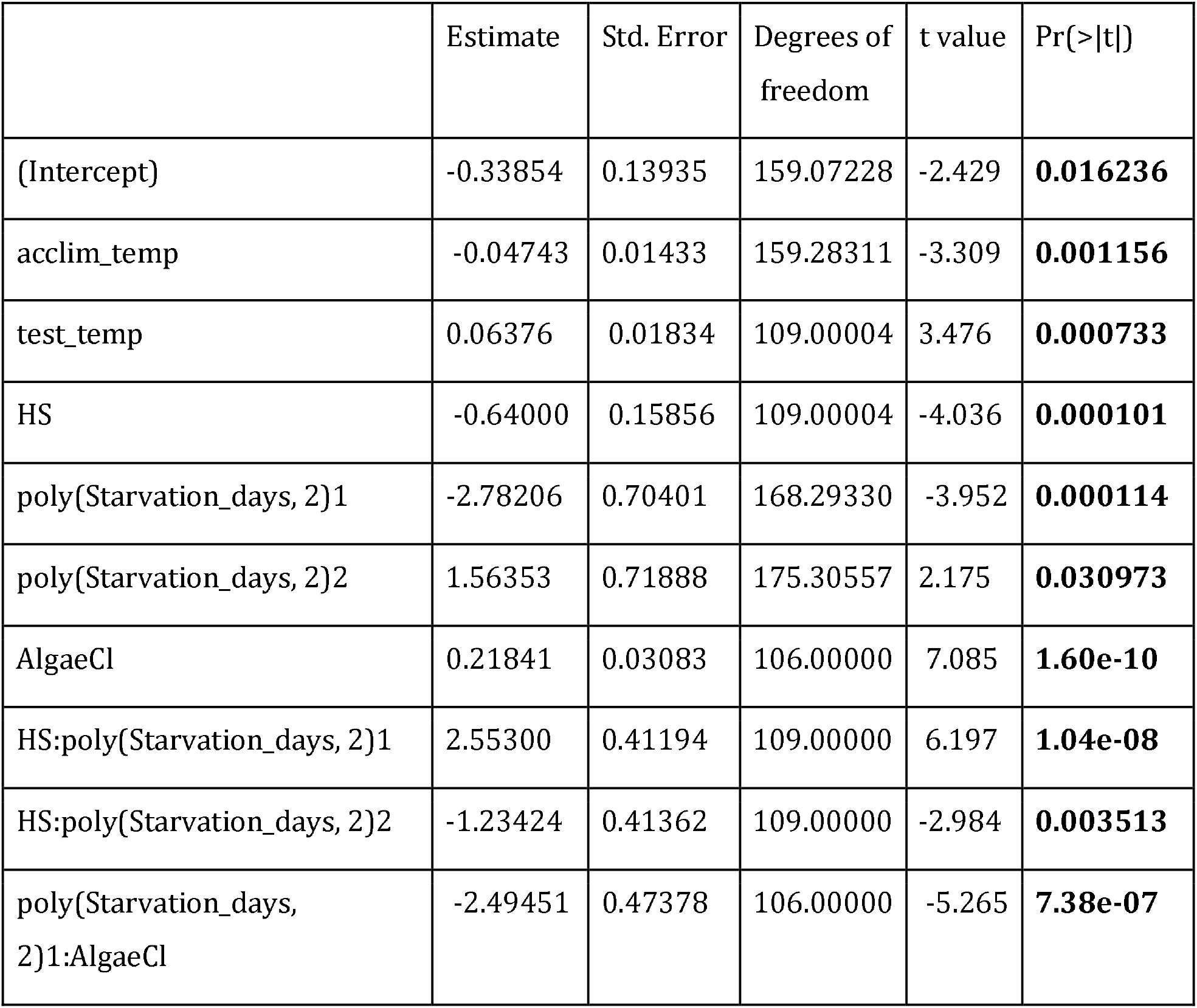
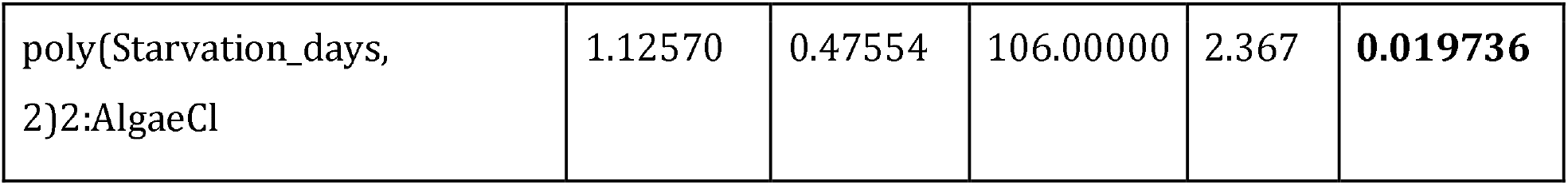
Summary table of the final model used in evaluating the effects of temperature, starvation and algal food source on oxygen uptake (Experiment 1).

**Table S4:** Summary table of the final model used in evaluating the effects of light intensity on oxygen uptake (Experiment 2).

|  | Estimate | Std. Error | t value | Pr(>\|t\|) |
| --- | --- | --- | --- | --- |
| (Intercept) | 0.496545 | 0.116198 | 4.273 | **3.86e-05** |
| Light intensityNL | -0.050163 | 0.038435 | -1.305 | 0.194317 |
| Heat shock | -0.746144 | 0.071168 | -10.484 | **< 2e-16** |
| Starvation_days | -0.011548 | 0.003295 | -3.505 | **0.000642** |
| HS:Starvation_days | 0.006968 | 0.002075 | 3.357 | **0.001052** |

### Supplementary Figures

Figure S1. The positive relationship between CT_first_ and CT_last_ (r ^2^=0.16) for Experiment 1 (A) and Experiment 2 (B). The solid blue line is the correlation between CT_first_ and CT_last_ and gray shading depicts 95% confidence intervals. The solid black line indicates X=Y (a perfect positive correlation).

Figure S2. A correlation matrix used to visualize correlations between the predictor variables we assessed in Experiment 1. A similar matrix could not be generated for Experiment 2 since the fixed effect “light intensity” only contained two levels (high light and normal light).

Figure S3. The mass in milligrams (mg) for specimens after the acclimation period where they were fed either *B. plumosa* (green violins) or *C.* cf. *linum* (blue violins) at each of the experimental temperatures. Slugs exposed to 17°C are indicated by violins outlined in orange and orange points and specimens exposed to 22°C have red outlines and red points. Only specimens exposed to 100µmol m ^-2^ s^-1^ light are plotted and points are slightly jittered on the x-axis.

